# The fragile X mental retardation protein promotes adjustments in cocaine self-administration that preserve reinforcement level

**DOI:** 10.1101/2020.07.06.190421

**Authors:** Jessica L. Huebschman, Megan C. Davis, Catherina Tovar Pensa, Yuhong Guo, Laura N. Smith

**Author notes:** Correspondence, Texas A&M University Health Science Center, Medical Research & Education Building I, Department of Neuroscience and Experimental Therapeutics, 8447 Riverside Parkway, Bryan, TX 77807.

## Abstract

The fragile X mental retardation protein (FMRP), an RNA-binding protein, regulates cocaine-induced neuronal plasticity and is critical for the normal development of drug-induced locomotor sensitization, as well as reward-related learning in the conditioned place preference assay. However, it is unknown whether FMRP impacts behaviors that are used to more closely model substance use disorders. Utilizing an intravenous cocaine self-administration (IVSA) assay in *Fmr1* knockout (KO) and wild type (WT) littermate mice, we find that, despite normal acquisition and extinction learning, *Fmr1* KO mice fail to make a normal upward shift in responding during dose-response testing. Later, with access to the original acquisition dose under increasing schedules of reinforcement (FR1, FR3, FR5), *Fmr1* KO mice earn significantly fewer cocaine infusions than WT mice. Importantly, operant conditioning with a palatable food reinforcer does not show similar deficits, indicating that our results do not stem from broad learning or reward-related deficits in *Fmr1* KO mice. Additionally, we find an FMRP target, the activity-regulated cytoskeleton-associated protein (Arc), to be significantly reduced in *Fmr1* KO mouse synaptic fractions from the nucleus accumbens following cocaine IVSA. Overall, our findings suggest that FMRP facilitates adjustments in drug self-administration behavior that generally serve to preserve reinforcement level, and combined with our similar IVSA findings in *Arc* KO mice, suggest Arc as a target of FMRP to investigate in behavioral shifts that occur when drug availability is altered.

## Introduction

Drugs of abuse drive synaptic plasticity throughout the brain, and vulnerability for substance use disorders and addiction is increased by previous drug exposure. Studies examining the pursuit of drugs of abuse in rodent models show that drug experience reduces the threshold dose required for learning (Piazza *et al.*, 1989; Horger *et al.*, 1990; Piazza *et al.*, 1990; Horger *et al.*, 1992) and increases willingness to work in these tasks. Drug-induced synaptic plasticity likely underlies this effect, as interrupting dopaminergic connections in the striatal-nigrostriatal pathway, which provides information from the ventral to the dorsal striatum, ultimately prevents the expression of habitual cocaine-seeking behavior without impairing new reward learning (Belin & Everitt, 2008). Further, AMPA receptor and dendritic spine upregulation following drug exposure and withdrawal appear to be critical for incentive sensitization, as measured by the ability of repeated cocaine treatment to increase later low-dose cocaine self-administration (Wang *et al.*, 2013), implicating drug-induced synaptic plasticity in the development of these maladaptive behaviors.

The fragile X mental retardation protein (FMRP) is an RNA-binding protein that regulates the transport, stability, and translation of hundreds of brain RNAs, playing a critical role in local translation and synaptic function (Darnell *et al.*, 2011). Loss of FMRP expression, as observed in fragile X syndrome (FXS), results in altered structural and functional synaptic plasticity, including widely reported changes in dendritic spine density (Irwin *et al.*, 2001; Nimchinsky *et al.*, 2001; Antar *et al.*, 2006; Grossman *et al.*, 2006; Neuhofer *et al.*, 2015), enhanced metabotropic glutamate receptor 5 (mGluR5)-dependent hippocampal long-term depression (Huber *et al.*, 2002), and impaired dopamine receptor D1 (DRD1)-mediated changes in cortical glutamate receptor 1 (GluR1) surface expression (Wang *et al.*, 2008). In the striatum, lack of FMRP has been associated with impaired N-methyl-D-aspartate receptor (NMDAR)-dependent long-term potentiation (LTP) and increased filopodia-type spines (nucleus accumbens; NAc, core region) (Neuhofer *et al.*, 2015), impaired striatal DRD1-receptor signaling (Wang *et al.*, 2008), and deficits in presynaptic function and stubby spine types (NAc, shell region) (Smith *et al.*, 2014).

FMRP has also been implicated as a regulator of drug-induced synaptic plasticity. *Fmr1* knockout (KO) mice, which lack FMRP, show increased dendritic spine density and synaptic strength in the NAc (shell) following repeated cocaine exposure at a time point when net changes in wild type (WT) animals are not observed (Smith *et al.*, 2014). Further, likely via its function in regulating synaptic plasticity, FMRP also facilitates drug-related behaviors, proving necessary for the normal development of locomotor sensitization following repeated cocaine administration, and while not required for all appetitive learning, is essential for drug-related reward learning using a low-moderate dose of cocaine in the conditioned place preference (CPP) assay (Smith *et al.*, 2014). Though presence of FMRP is clearly important for the effects of cocaine on synaptic structure and function, as well as for drug-related behavioral alterations, it is unknown if its function influences vulnerability to other addiction-related behaviors.

Here, based on prior studies, we hypothesized that absence of FMRP would impair cocaine-, but not food-reinforced operant responding. We have recently reported impaired cocaine intravenous self-administration behavior (IVSA) in a KO mouse lacking the activity-regulated cytoskeleton-associated protein (Arc), a target of FMRP. Therefore, we also planned examinations of protein expression levels for Arc and two additional downstream targets of FMRP in cellular fractions following contingent cocaine exposure in the presence and absence of FMRP regulation. Based on our prior observations, we expected striatal Arc protein levels to be significantly decreased in cellular fractions from NAc tissue in the absence of FMRP.

## Methods

### Animals and drugs

WT and *Fmr1* KO littermates were produced at the Texas A&M University Health Science Center by *Fmr1*^−/y^ and *Fmr1*^−/+^ breeders (Jackson Laboratory, Stock # 003025) that have been maintained on a C57BL/6N (Charles River) background for more than 25 generations, with regular backcrossing. Offspring were group-housed by sex under standard laboratory conditions in ventilated cages with a 12-hour light/dark cycle (on at 0600) and *ad libitum* access to standard mouse chow and water. A total of 44 naïve adult male mice (*Fmr1*^−/y^ and *Fmr1*^−/+^; 10-15 weeks at testing) were designated for either food or drug operant conditioning experiments. Animals were not food-deprived, and cocaine IVSA mice were not food trained. Cocaine intravenous solutions were prepared with (-)-cocaine hydrochloride (NIDA Drug Supply Program; cat. #9041-001) in 0.9% USP-grade saline and sterile-filtered (0.02 μm). All procedures were conducted in compliance with the Texas A&M University Institutional Animal Care and Use Committee.

### Jugular Vein Catheter Implantation Surgery

Indwelling back-mount catheters constructed in-house using commercial intravenous jugular catheters (SAI Infusion Technologies) were implanted under oxygen/sevoflurane vapor, following a previously described protocol (Thomsen & Caine, 2005). Briefly, anchored catheters extended 1.0-1.2 cm into the jugular vein and passed subcutaneously over the same shoulder to the base seated above the mid-scapular region. When placement into the right jugular vein was unsuccessful or a replacement catheter was necessary, the left vein was used. Following surgery, mice were given one day without handling before daily administration of antibiotic/anti-clotting solution (30 USP units/ml heparin and 67 mg/mL cefazolin in 0.02 mL of 0.9% saline) during the rest of the 7-day recovery period. Thereafter, catheters were flushed with 0.03 μL of heparinized (30 USP U/mL) saline (0.9%) before and after self-administration sessions. Patency was evaluated periodically throughout and at the end of the experiment by flushing 0.03 mL of 15 mg/mL ketamine/0.75 mg/mL midazolam through the catheter; loss of righting reflex within 3 seconds was required.

### Operant Conditioning

Operant chambers were each housed in a light- and sound-attenuating chamber and contained two nose-poke ports (active and inactive, randomly assigned), fixed on either side of a liquid food delivery magazine, with a cue light above each. Mice were trained to self-administer either a liquid food reward (20 μL/administration Ensure^®^ nutritional drink, vanilla flavor, 100%; Abbott Laboratories) delivered via a dipper cup or an intravenous infusion of cocaine (1.0 mg/kg/infusion in sterile 0.9% (w/v) NaCl) delivered via a syringe pump. Drug infusion volume was 0.56 mL/kg, and concentration was adjusted according to the desired dose (e.g., for a 32 g mouse, each infusion equaled 18 μL, and to achieve a dose of 1.0 mg/kg/infusion, a concentration of 1.8 mg/mL was used). The length of each infusion was calculated as weight (kg)/0.01 s (i.e., a 32 g mouse receives 3.2 s infusions). Both experiments ran 5 days/week. Food operant sessions were 2 hrs per day and terminated early if 100 reinforcers were reached. IVSA sessions were 3 hrs per day and terminated early at different reinforcer limits, depending on dose (1.0 mg/kg/inf = 30 max; all lower doses = 100 max; 3.2 mg/kg/inf = 10 max with 10 min timeouts if ≥ 2 reinforcers in ≤ 10 min). A house light signaled reinforcer availability, which was turned off following each delivery (“time-out”; 10 s for food, 20 s for drug). Entries into all ports and magazines were recorded throughout each session by beam break. Rarely, we found that blockage of a beam detector falsely registered as a very high beam break value; therefore, measurements of 125 or greater in a single session were excluded from analysis and presentation.

Experimental phases ran as shown in Figure 1. Acquisition sessions began with illumination of the active port cue light and either availability of liquid reinforcer for 60 s (food) or a single infusion to prime the jugular catheter (drug). Nose-pokes in the active port resulted in reinforcement and simultaneous illumination of the cue light on a fixed-ratio 1 (FR1) schedule of reinforcement, while nose-pokes in the inactive port had no consequence. Sessions for all other test phases were conducted exactly the same, except that during extinction sessions active nose-pokes resulted in either water delivery (food) or no consequence (drug). Each mouse’s acquisition and extinction phases were ended upon completion of pre-defined learning criteria. Acquisition criteria were defined as two consecutive sessions having ≥20 (food) or ≥15 (drug) reinforcers with no more than 20% variation in response number between sessions, and at least 70% discrimination for the active port. Overnight sessions, otherwise identical to acquisition sessions, were given after 5 and 10 days to animals that had not yet reached acquisition criteria, with a similar number from each group requiring both overnight sessions to meet criteria (drug: 0 WT, 3 *Fmr1* KO; food: 1 WT, 2 *Fmr1* KO). Overnight sessions are not included in data analysis or graphs. Extinction sessions ran until active port responding dropped to a portion of the average of the last two days of acquisition: ≤ 50% for food operant (for two consecutive days) or ≤ 30% for drug IVSA (one day). Animals that did not meet extinction criteria within 15 days were dropped from the study (drug studies: 2/group; food studies: 0/group). Re-acquisition sessions were conducted until reinforcers earned in a single session returned to ≥20 (food) or ≥15 (drug). All animals met re-acquisition criteria in 1-2 sessions. For dose-response testing, a single concentration was made available each session over either two days (food; 0, 3, 10, 32, and 100% Ensure^®^, in water) or one (drug operant studies; 0, 0.01, 0.03, 0.1, 0.32, 1.0, and 3.2 mg/kg/infusion, in 0.9% saline). Doses were presented in sequential order, with the starting dose counterbalanced across groups using a Latin square design. For increasing cost testing in IVSA, mice were returned to the acquisition dose of cocaine (1.0 mg/kg/infusion), starting on an FR1 schedule of reinforcement, then were moved to FR3, followed by FR5 (two consecutive sessions each).

**Figure 1.**
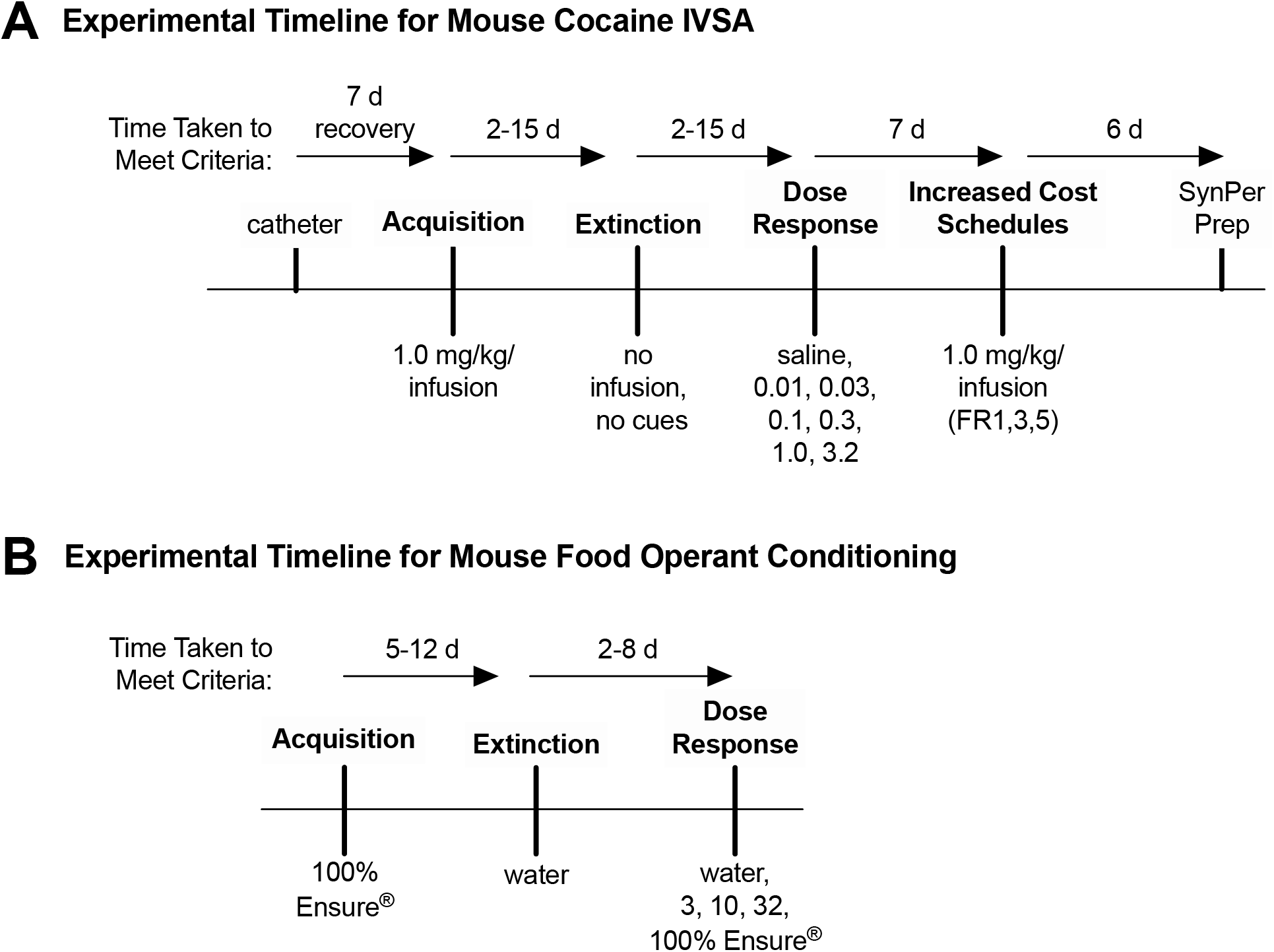
Experimental timeline for mouse IVSA (A) and food operant conditioning (B) studies.

### Synaptosome Isolation and Western Blotting

Synaptosomes were only prepared from mice completing all phases of IVSA. Following live decapitation, NAc (core + shell; starting Bregma ~1.65 mm) and dorsal striatum (DS; medial + lateral; starting Bregma ~1.42 mm) were rapidly dissected from coronal (1 mm) slices, using the anterior commissure as a guide. All tissue processing steps were performed at 4°C or on ice. Samples were lysed in Syn-PER Synaptic Protein Extraction Reagent (ThermoFisher 87793) containing 1X cOmplete EDTA-free Protease Inhibitor Cocktail (Roche) using a handheld pellet pestle mixer and centrifuged at 1200 × *g* for 10 min. After reserving a portion of the resulting supernatant (homogenate), the remainder was centrifuged for 20 min at 15,000 × *g*. The second supernatant (cytosolic fraction) was saved, and the remaining pellet (synaptosome) was resuspended in Syn-PER, according to manufacturer directions. Protein quantification was performed on all fractions using the DC Protein Assay Kit (BioRad). Samples (40 μg total protein/well) were electrophoresed on 8% sodium dodecyl sulfate-polyacrylamide (SDS-PAGE) gels and transferred (30 V; Thermo-Fisher mini-blot wet transfer system; 1 hr) to Immobilon FL polyvinylidene fluoride (PVDF) membranes (Millipore) for immunoblotting. Membranes were rinsed (mQH_2_O), then blocked in casein buffer (LI-COR 927-40200; shaking at room temperature; 1 hr) and incubated (shaking at 4°C; overnight) in primary antibody diluted in casein buffer. Primary antibodies were ARC (Synaptic Systems 156-002, 1:2000), PSD-95 (Millipore Sigma MABN68, 1:1000), NR2B (Cell Signaling 14544S, 1:1000), and GAPDH (Fisher-Scientific AM4300, 1:1000). Afterward, membranes were rinsed in 1X TBS-T and incubated (shaking at room temperature; 1 hr) in a secondary antibody diluted in casein buffer plus 0.01% SDS and 0.2% Tween. Secondary antibodies were goat anti-rabbit or anti-mouse IgG polyclonal IRDye^®^-680RD or IRDye^®^ 800CW (1:20,000; LI-COR). After final rinses in 1X TBS-T, membranes were scanned and analyzed using Odyssey CLX and ImageStudio (LI-COR).

### Statistical Analysis

Statistical analysis and results are listed in Table S1. For behavior testing, responses on all ports were initially analyzed with two- (genotype × port), threeÒ (genotype × port × session), or four-way (genotype × port × schedule × day) ANOVAs, having both between- (genotype) and within-subjects, or repeated measures (RM), factors (port, session, schedule, day). For western blots, relative expression of the protein of interest was first normalized to a housekeeping protein (GAPDH) and then to the WT S1 fraction before analysis with two-way (genotype × fraction) ANOVA. GAPDH was chosen as a normalization protein because it is not suspected to be an FMRP target (Darnell *et al.*, 2011). Standard error was propagated to account for normalization steps. Significant interactions were followed by additional ANOVAS (i.e., one-, two-way, etc.), paired t-tests, and/or Bonferroni or Tukey post hoc analyses, as appropriate, to determine simple main effects (SMEs). When Mauchley's test of sphericity was significant, either Greenhouse-Geisser (G-G; when Epsilon <0.75) or Huynh-Feldt (H-F; when Epsilon>0.75) corrections were used (see Table S1). Comparison of group survival distributions for reaching criteria were performed for acquisition and extinction phases of food operant and IVSA assays using Kaplan-Meier log rank tests (Mantel-Cox). Mice that failed to reach criteria were included in survival analyses, but are not shown in any graphs. All statistics were performed using GraphPad Prism, except SPSS software was used to handle complex data sets/analyses (e.g., missing values, three-way, four-way, and MV ANOVAs). Significance was set at alpha = 0.05.

## Results

### FMRP is not required for acquisition of cocaine or food reward-related operant conditioning

To determine whether FMRP is necessary for reward-related operant learning, we measured the acquisition of cocaine (intravenous; Fig. 2A) and food (Ensure^®^ liquid reinforcer; Fig. 2D) self-administration in separate cohorts of *Fmr1* KO and WT littermates. To limit contributions of overtraining to group differences, acquisition and extinction phases during each test were ran to a set of criteria selected to indicate learning (see Methods); thus, individual animals differed in the number of sessions during each phase (Fig. 1; Figs. S1, S2). Log rank tests showed that survival distributions of *Fmr1* KO and WT mice for reaching acquisition criteria were not significantly different for either drug (Fig. 2B) or food (Fig. 2E) reinforcers, and similar numbers from each group failed to meet criteria (drug: 6 WT, 2 *Fmr1* KO; food: 1 WT, 0 *Fmr1* KO). Comparing the first and final acquisition days of drug IVSA (Fig. 2C) and separately, of food operant conditioning (Fig. 2F) revealed no main effects or interactions involving genotype. Port x session interactions were significant for drug- (*F(1, 24) = 72.277, p < 0.0001)* and food-reinforced (*F(1, 18) = 104.585, p < 0.0001)* acquisition phases. Follow-up analyses for the cocaine IVSA study showed a simple main effect (SME) of port at the last (*F(1, 26) = 499.497, p < 0.0001)*, but not first, session, while there were significant SMEs of session for both the active (*F(1, 26) = 23.159, p < 0.0001)* and inactive (*F(1, 25) = 25.915, p < 0.0001)* ports. For the food study there were significant SMEs of port at both first (*F(1, 19) = 5.438, p = 0.031)* and last (*F(1, 19) = 206.640, p < 0.0001)* sessions, while SMEs of session reached significance for the active port (*F(1, 19) = 172.929, p < 0.0001)* and trended toward significance for the inactive port (*F(1, 19) = 4.279, p = 0.053)*. These results are consistent with normal acquisition of both food- and drug-reinforced operant conditioning across groups, suggesting that learning of these tasks is not impacted by loss of FMRP expression.

**Figure 2.**
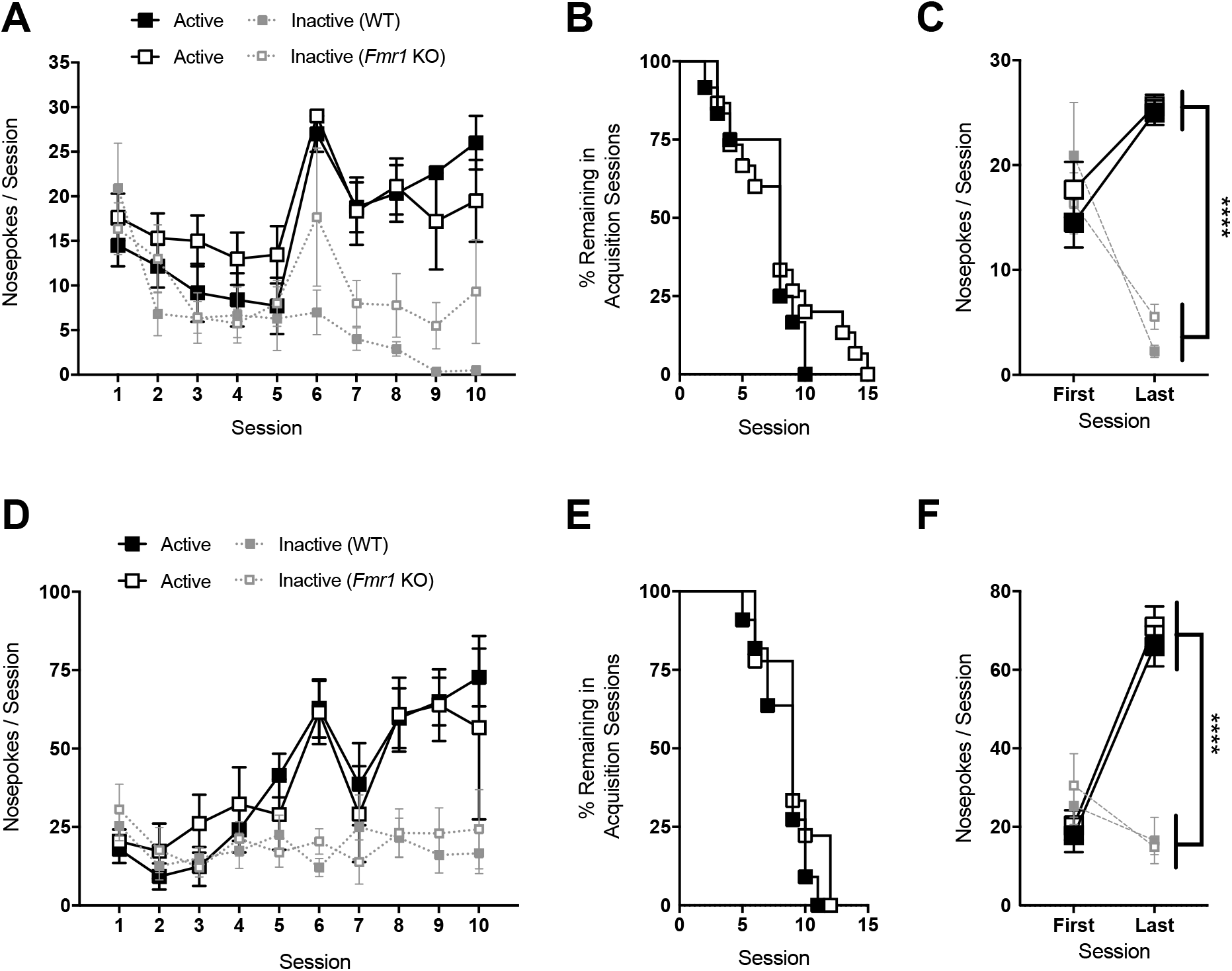
Average nose-pokes (active port, large boxes; inactive port, small boxes), excluding responses made during time-out periods, for wild type (WT) and *Fmr1* knockout (KO) mice during acquisition sessions for drug (A; 1.0 mg/kg/infusion, cocaine; *n* = 12 WT, 15 KO) and food (D; 100% vanilla-flavored Ensure^®^; *n* = 11 WT, 9 KO) studies. Only sessions where > 25% of animals per group remained in the respective phase of behavioral testing are shown. The rate at which animals met acquisition criteria in each group for drug (B) and food (E) studies was not different. Animals that did not reach criteria are not included in figures, but were included in group survival analysis. As animals met criteria they moved on to the next phase of testing, so number of acquisitions sessions varies by animal. There was no effect of genotype on average nose-pokes for first and last sessions of acquisition for either drug (C) or food (F) studies. ****p < 0.0001; data shown are mean ± S.E.M.

### FMRP is not required for extinction of cocaine or food reward-related operant conditioning

To test whether FMRP has a role in the extinction of reward self-administration, we compared the responses of *Fmr1* KO and WT animals following successful acquisition under conditions where active port activity no longer resulted in the delivery of a reinforcer (Figs. 3A, 3D). There were no significant differences between the genotypes in survival distributions for meeting extinction criteria in either drug (Fig. 3B) or food (Fig. 3E) experiments. Similar numbers of animals per group failed to extinguish reward-seeking behavior within 15 sessions (drug: 5 WT, 4 *Fmr1* KO; food: 0 WT, 0 *Fmr1* KO) and were removed from further analysis. Comparing the first and last cocaine extinction sessions (Fig. 3C), we observed a significant port × session interaction (*F(1, 21) = 22.462, p < 0.0001)*, while no interactions or main effects involving genotype were observed. Follow-up to this interaction showed a significant SME of session for both the active (*F(1, 22) = 52.214, p < 0.0001*) and inactive (*F(1, 25) = 10.466, p = 0.003*) ports between the first and last day. There was also a significant effect of port on the first extinction session (*F(1, 22) = 16.352, p = 0.001*). In extinction of food-operant conditioning (Fig. 3F), the overall ANOVA similarly showed a significant port x session interaction (*F(1, 17) = 37.790, p < 0.0001)*, with no main effects or interactions involving genotype. Follow-up analyses showed significant SME of port at both the first (*F(1, 18) = 49.484, p < 0.0001)* and last (*F(1, 19) = 11.978, p = 0.003)* sessions. Significant session SMEs were present for the active port (*F(1, 18) = 40.236, p < 0.0001)*, but not the inactive port.

**Figure 3.**
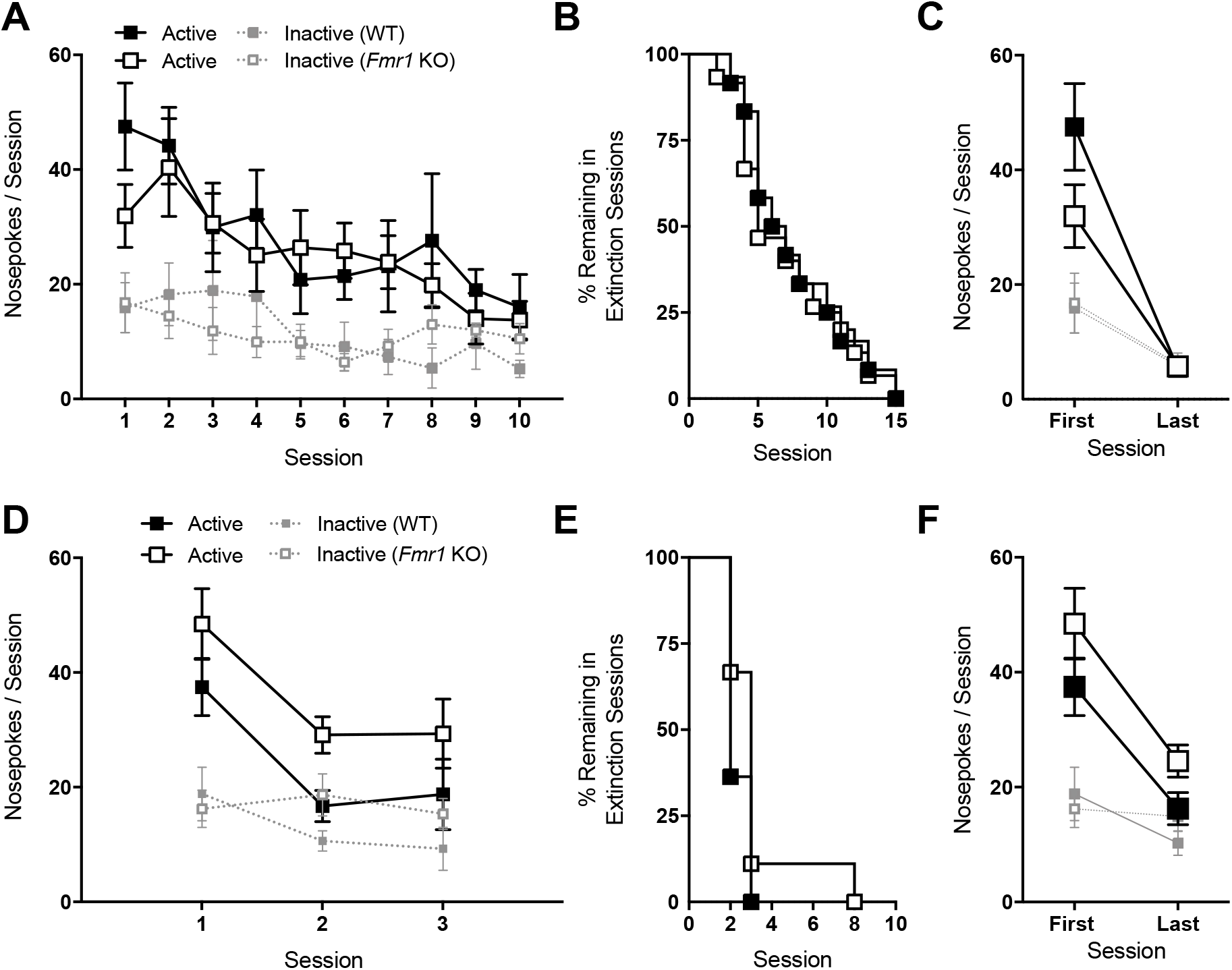
Average nose-pokes (active port, large boxes; inactive port, small boxes) for wild type (WT) and *Fmr1* knockout (KO) mice during extinction sessions for drug (A; *n* = 12 WT, 15 KO) and food (D; *n* = 11 WT, 9 KO) studies. Only sessions where > 25% of animals per group remained in the respective phase of behavioral testing are shown. The rate at which animals met extinction criteria in each group for drug (B) and food (E) studies was not different. Animals that did not reach criteria are not included in figures, but were included in group survival analysis. As animals met criteria they moved on to the next phase of testing, so number of extinction sessions varies by animal. There was no effect of genotype on average nose-pokes for first and last sessions of extinction for either drug (C) or food (F) studies. Data shown are mean ± S.E.M.

### FMRP is required for a normal dose-response in cocaine, but not food, operant conditioning

We next evaluated self-administration behavior for both drug and food reinforcers across a range of doses. For cocaine experiments, there was a significant three-way interaction between genotype, dose, and port (Fig. 4A; *F(2.2, 36.8) = 3.235, p = 0.047*). The follow-up two-way ANOVA for the active port alone revealed a significant interaction of genotype and dose (*F(2.1, 39.6) = 4.873, p = 0.012*), while there were no significant interactions or main effects when the same analysis was performed for inactive port responses. When WT animals were analyzed alone, there was a significant port x dose interaction (*F(2.1, 12.7) = 9.636, p = 0.003*), and follow up one-way RM ANOVAs showed a significant SME of port at the 0.1, 0.32, and 1.0 mg/kg/infusion doses *(p = 0.026, 0.00014, < 0.0001, respectively)*. One-way ANOVA over unit dose for WT animals on the active port showed a significant main effect of dose (*F(2.1, 16.6) = 14.902, p < 0.0001*), with responses for the 0.32 unit dose significantly different from those for the 0.01, 0.032, 1.0, and 3.2 unit doses (Bonferroni pairwise comparisons, *p = 0.01, 0.014, 0.009, 0.002*, respectively), and the 1.0 unit dose responses significantly different than those for the 3.2 unit dose (*p < 0.0001*; see Table S1). A two-way ANOVA for KO mice alone also showed a significant dose × port interaction (*F(2.0, 22.4) = 4.703, p = 0.019*) and follow up one-way ANOVAs revealed a significant SME of port at the 0.1, 0.32, and 1.0 mg/kg/infusion doses. One-way ANOVA over unit dose for KO animals on the active port showed a significant dose main effect (*F(2.0, 21.9) = 4.088, p = 0.031*), with a significant difference in responses between the 1.0 and 3.2 unit doses *(Bonferroni pairwise comparison, p < 0.0001)*. Direct comparisons of WT and KO active port responses at each dose individually revealed a significant simple main effect of genotype at the 0.32-unit dose, with WT mice earning significantly more infusions than KO at this dose (*F(1,17) = 8.023, p = 0.011*).

**Figure 4.**
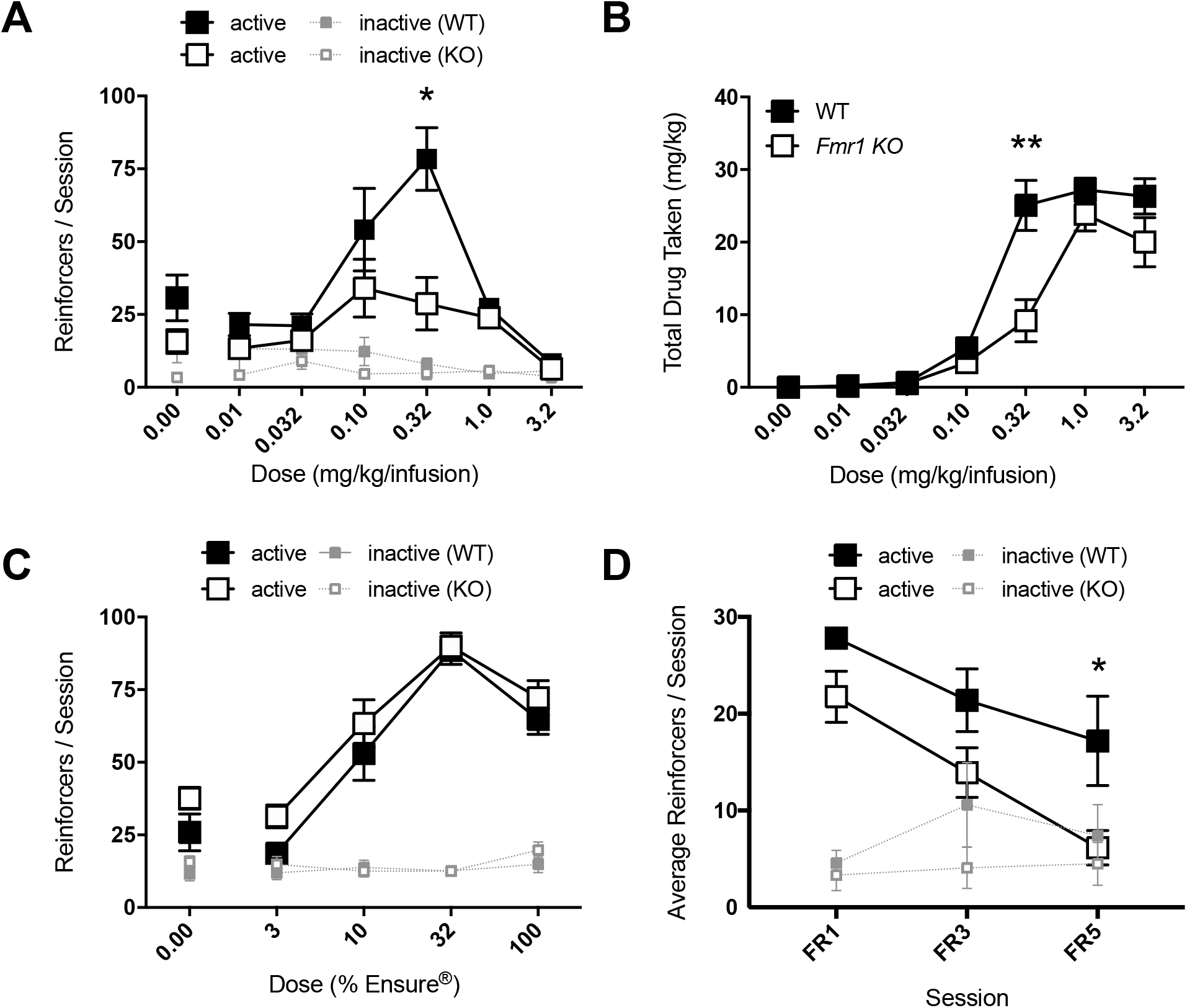
Average nose-pokes (active port, large boxes; inactive port, small boxes), excluding those made during time-out periods, for wild type (WT) and *Fmr1* knockout (KO) mice during IVSA dose-response testing (A; *n* = 9 WT, 12 KO). KO mice earn significantly fewer reinforcers than WT mice at the 0.32 mg/kg/infusion dose, resulting in lower total drug intake at this dose (B). There were no differences between genotypes during dose-response testing for a food reinforcer (C; *n* = 11 WT, 9 KO). (D) Average nose-pokes for WT and *Fmr1* KO mice under increasing schedules of reinforcement (FR1, FR3, FR5; *n* = 5 WT, 6 KO). ^#^p < 0.1, *p < 0.05, **p < 0.01; data shown are mean ± S.E.M.

When we compared total drug intake across the dose-response test, we also observed a significant interaction between genotype and dose (Fig. 4B; *F(2.7, 52.2) = 5.374, p = 0.003*), with *Fmr1* KO mice receiving significantly less cocaine than their WT counterparts at the 0.32-unit dose (MV ANOVA, *F(1, 19) = 12.526, p = 0.002)*. One-way ANOVA for the WT group showed a significant SME of unit dose on drug intake (*F(2.2, 17.5) = 60.58, p < 0.0001)*, with Bonferroni pairwise comparisons showing significant differences between each of the lowest three and highest three doses (0.32 > 0.01, 0.032, 0.1; *p = 0.001, 0.002, 0.003*; 1.0 > 0.01, 0.032, 0.1; *p < 0.0001, 0.0001, 0.0001*; 3.2 > 0.01, 0.032, 0.1; *p ≤ 0.0001, 0.0001, 0.001*; see Table S1). For KO mice there was a significant SME of unit dose for drug intake (*F(2.5, 27.4) = 31.402, p < 0.0001)*, and Bonferroni pairwise comparisons showed that at the 1.0-unit dose, KO mice had higher intake than at each of the lower doses (1.0 > 0.01, 0.032, 0.1, and 0.32; *p ≤ 0.0001, 0.0001, 0.0001, 0.004*, respectively), more at the 3.2-unit dose than at the lowest three doses (3.2 > 0.01, 0.032, 0.1; *p = 0.002, 0.002, 0.008*, respectively), and significantly more at the 0.032-unit dose than the 0.01-unit dose (*p = 0.016*).

Three-way ANOVA for reinforcers earned during food study dose-response experiments showed a significant port x concentration interaction (*F(3, 54) = 79.627, p < 0.0001)*, while there was no interaction or main effect involving genotype (Fig. 4C). We observed significant SMEs of concentration on the active (*F(3, 54) = 73.575, p < 0.0001*) and inactive (*F(3, 54) = 3.159, p = 0.03*) ports. We investigated each genotype separately to determine whether operant conditioning learning deficits are present in *Fmr1* KO mice. There were port × concentration interactions for each group (WT: *F(3, 30) = 40.546, p < 0.0001*; KO: *F(3, 24) = 45.959, p < 0.0001)*. In addition, there were SMEs of concentration on the active port for both WT mice (*F(3, 30) = 42.305, p < 0.0001)* and KO (*F(3, 24) = 33.915, p < 0.0001*) groups, with significant Bonferroni pairwise comparisons between all doses except 30% and 100% for both the WT and KO groups separately (*see Table S1*). KO mice also showed a significant SME of concentration on the inactive port (*F(3, 24) = 4.479, p = 0.012*); however, none of their responses on this port differed significantly between concentrations.

Finally, we analyzed WT and *Fmr1* KO mouse behavior on an increasing FR schedule of reinforcement, with each schedule being presented on two consecutive days before proceeding to the next *(i.e. FR1, FR1, FR3, FR3, FR5, FR5*; Fig. 4D). Four-Way ANOVA (genotype × port × schedule × day) showed significant interaction between port and schedule (*F(2,18) = 28.997, p < 0.0001*) and a main effect of genotype (*F(1,9) = 5.795, p = 0.039*). Follow-up analyses showed a significant port effect at FR1 (*F(1, 10) = 117.356, p < 0.0001*) and FR3 (*F(1, 10) = 15.121, p = 0.003*), but not FR5. Looking at active port responses alone, there was a significant effect of schedule (*F(1.6, 15.9) = 25.741, p < 0.0001*) with Bonferroni pairwise comparison revealing a significant decrease in responding at this port from FR1 to FR3 (*p = 0.003*) and FR3 to FR5 (*p = 0.007*). There was no effect of schedule on inactive port responses, indicating lack of port effect at FR5 is a result of decreased responding at the active port, likely due to the increased difficulty of the task rather than a change in port preference. To better interpret the overall main effect of genotype given the interaction of port and schedule described above, we conducted additional MANOVA analysis for each port at FR1, 3, and 5 and found a significant effect of genotype in active port responses at FR5 (*F(1, 10) = 5.739, p = 0.04*), with KO animals receiving significantly fewer reinforcers at this schedule of reinforcement.

### FMRP promotes expression of ARC in the nucleus accumbens following cocaine exposure

Based on our recent findings that the FMRP target, Arc, is also critical for vertical shifts in the cocaine IVSA dose-response function (Penrod *et al.*, 2020), we examined ARC protein expression in WT and *Fmr1* KO animals in cell fractions (S1: total homogenate, S2: cytosolic, P: synaptosome) isolated from NAc and DS one day following their last cocaine self-administration session. For comparison, we also evaluated the expression of two other FMRP targets, PSD-95 and NR2B. For ARC, there was a significant interaction in NAc between genotype and cell fraction (*F(2,18) = 6.228, p =0.0088*), with post-hoc analysis revealing a significant deficit in ARC expression in the synaptosome (P fraction) of *Fmr1* KO mice (Fig. 5D; *p = 0.0039*). The expression of ARC across cell fractions in the DS appears similar to those observed in the NAc (Fig. 5H); however, analysis of expression levels in DS showed a significant main effect of cell fraction only (*F(2,18) = 10.31, p = 0.0010)*. In contrast to ARC results in the NAc, only a significant main effect of fraction was observed for PSD-95 expression in both NAc (Fig. 5B; *F(2,18) = 40.17, p < 0.0001*) and DS (Fig. 5F; *F(2,18) = 16.04, p < 0.0001*). Similarly, we observed a main effect of fraction for NR2B expression in both NAc (Fig. 5C; *F(2,18) = 17.49, p < 0.0001*) and DS (Fig. 5G; *F(2,18) = 11.76, p =0.0005*).

**Figure 5.**
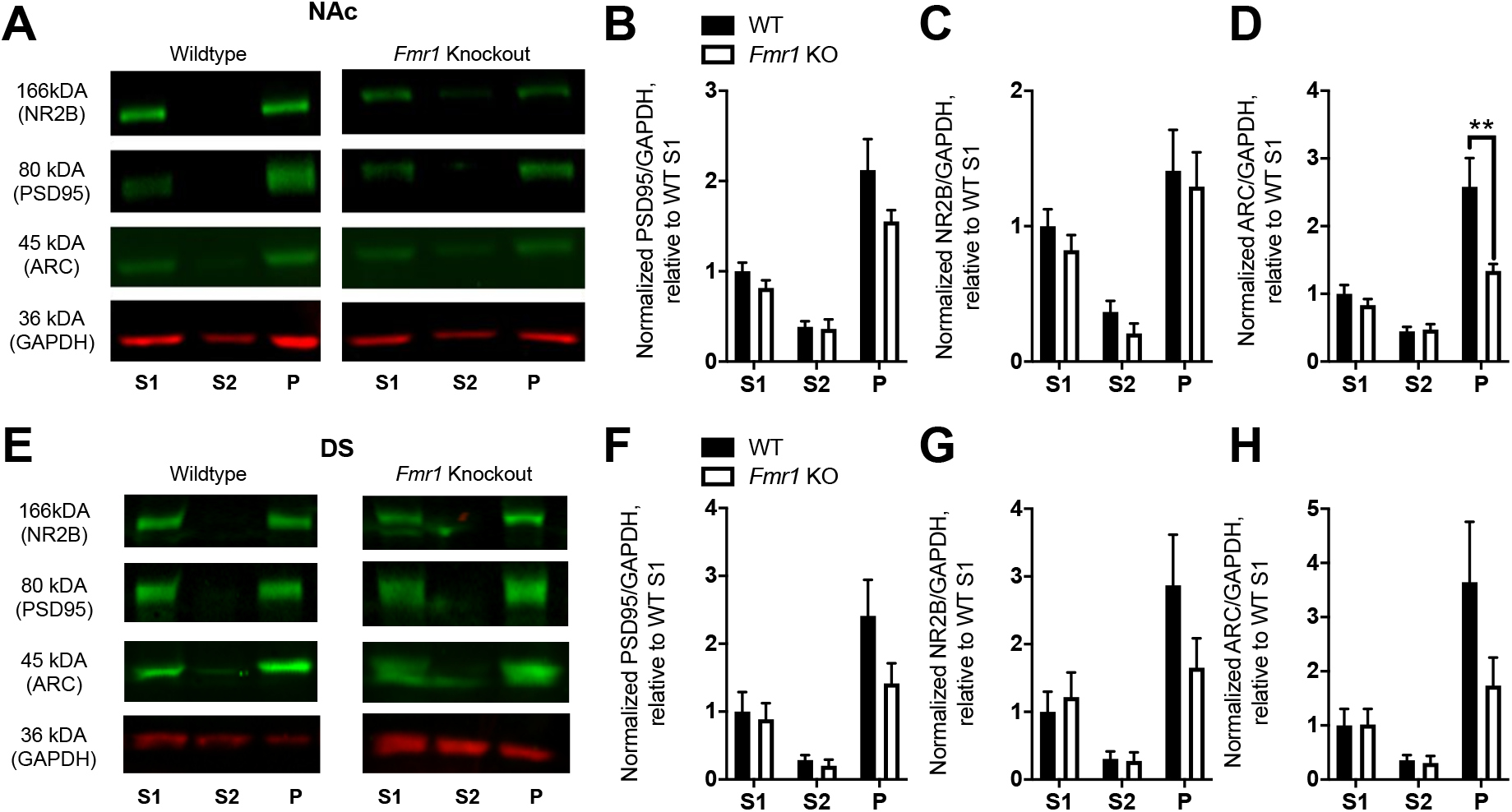
Expression of FMRP targets (PSD-95, NR2B, ARC) in cell fractions (S1: total homogenate, S2: cytosolic, P: synaptosome) from the nucleus accumbens (NAc; A-D) and dorsal striatum (DS; E-H) of wild type (WT) and *Fmr1* knockout (KO) mice ~ 24 hours after the final cocaine self-administration session (*n* = 4 WT, 4 KO). Expression values for each protein were normalized to the housekeeping protein GAPDH and to the WT S1 fraction. (A, E) Representative blots, all bands shown for each region are from a single blot. *Fmr1* KO animals show significantly decreased expression of ARC in NAc synaptosome fraction compared to WT animals. ** p < 0.01; data shown are mean ± S.E.M.

## Discussion

Drugs of abuse, like cocaine, induce synaptic plasticity throughout the striatum, a process thought to contribute to changes in the incentive salience of drug-related cues and experience, as well as to the perceived hedonic value of drugs. FMRP, an RNA-binding protein that mediates synapse plasticity, regulates cocaine-induced structural changes in the NAc, and its absence significantly weakens cocaine-related behavioral responses (Smith *et al.*, 2014). In this introductory investigation of FMRP’s role in drug self-administration, using a total knockout mouse, we show that FMRP also facilitates behavioral plasticity associated with operant cocaine self-administration. Mice lacking FMRP do not maintain normal drug-taking behavior, an effect observed post-extinction in both their dose-response curve and following a return to normal task requirements and acquisition dose under increasing infusion cost. Deficits do not extend to operant responding for a natural reward, indicating normal learning ability in KO mice. Following cocaine IVSA, protein expression of Arc, but not NR2b or PSD-95, is significantly reduced in NAc synaptic cellular fractions from *Fmr1* KO, compared to WT, mice.

The most striking finding here is a significantly flattened cocaine dose-effect function observed in *Fmr1* KO mice, in contrast to WT mice. Drugs of abuse generally support inverted U-shaped dose-response curves in the IVSA task, as rodents adjust response frequency to maintain a targeted minimum satiety according to the unit dose (Tsibulsky & Norman, 1999) and other factors (Panlilio *et al.*, 2003). Left or right shifts in the dose-response curve can indicate changes in sensitivity or tolerance to reinforcing and/or rate-limiting effects of the drug (Schenk & Partridge, 1997). Since average *Fmr1* KO responding remains at or below WT at every dose, notwithstanding occlusion by disparities in rate-limiting effects of cocaine, we can be reasonably sure that differences in sensitivity or tolerance to the reinforcing effects of cocaine do not explain our findings. Instead, the deficit we observe in *Fmr1* KO mice, with a significant group difference at the WT peak responding dose, is consistent with a downward shift.

Vertical-shifted dose-response functions have been primarily discussed in terms of altered hedonic experience and/or set point, with prior drug experience and escalation of voluntary intake being associated with upward shifts respective to the normal inverted curve (Ahmed & Koob, 1998). While vertical shifts cannot be used as evidence of incentive sensitization, they are not inconsistent with such interpretations (Robinson & Berridge, 2003). Upward shifts are also observed in rodents with high locomotor responses to novel contexts. These “high-responders” display riskier drug-related behaviors overall, including self-administration of lower drug doses and stronger responding on increased schedules of reinforcement (Piazza *et al.*, 2000; Deroche-Gamonet *et al.*, 2004; Kasanetz *et al.*, 2010). Similarly, when we return to the acquisition phase dose (1.0 mg/kg/infusion) and task requirements (FR1) after dose-response testing and then increase response requirements (FR3, FR5), we observe that WT mice, as a group, earn more reinforcers than *Fmr1* KO mice, and also work harder than KO mice for cocaine at the highest cost requirement. Together, these findings suggest that FMRP promotes adjustments in responding for cocaine in the IVSA task, particularly those requiring increased effort, ultimately supporting the preservation of reinforcement level.

Flattened dose-response curves following normal acquisition in the cocaine IVSA assay are not commonly observed, though there are examples (Deroche-Gamonet *et al.*, 2003; Thomsen *et al.*, 2009). We recently reported this effect in mice lacking the immediate early gene, Arc (Penrod *et al.*, 2020), a target of FMRP (Zalfa *et al.*, 2003). Like *Fmr1* KO mice, this flattened curve in *Arc* KO mice is also observed following normal task acquisition and extinction. In both studies, major group differences were only observed after repeated cocaine exposures (acquisition) and drug abstinence (extinction), so it is tempting to speculate that one or both experiences are responsible for later observations. Intriguingly, despite similar profiles in cocaine IVSA, conditioned place preference and particularly locomotor sensitization to cocaine in these two KO mouse groups markedly contrast with one another (Smith *et al.*, 2014; Smith *et al.*, 2016; Salery *et al.*, 2017; Penrod *et al.*, 2020). There are also subtler, yet notable, differences in our cocaine IVSA observations between *Fmr1* KO and *Arc* KO mice. During dose-response testing, in contrast to the *Fmr1* KO deficit at the WT peak dose, *Arc* KO mice deficits covered most of the descending limb of the curve. Later, under increasing cost schedules, *Arc* KO and WT mice showed no signs of diverging at any cost level (FR1, FR2, FR3; note FR5 was not included), while *Fmr1* KO mice showed fewer responses than WT across the tested schedules (FR1, FR3, FR5). In part, these results are indicative of disparities in willingness to work for cocaine, with loss of FMRP resulting in a deficit not present in *Arc* KO mice. Potential relationships between either KO group’s non-contingent and contingent cocaine-induced behaviors, perhaps commonly influenced by sub-circuit level alterations, remain to be determined.

Arc is considered an effector immediate early gene, a reference to its direct, activity-induced influence on cellular function, namely removing glutamatergic AMPA receptors from synapses (Chowdhury *et al.*, 2006), and its expression (mRNA and protein) is induced by cocaine exposure (Fosnaugh *et al.*, 1995; Samaha *et al.*, 2004). Arc’s potential role in mediating long term molecular and behavioral responses to cocaine has also been widely reported (Hearing *et al.*, 2008; Zavala *et al.*, 2008; Freeman *et al.*, 2010; Salery *et al.*, 2017). Due to its status as a target of FMRP, we questioned whether dysregulation of Arc in *Fmr1* KO mice might contribute to similarities in IVSA behavior across these KO mouse lines. Since our observations have relevance for understanding early drug-related behavioral regulation, and the NAc has been linked to initial drug reward, we hypothesized that *Fmr1* KO mice would show Arc protein expression deficits in cellular compartments for this brain region. Based on our results in total *Arc* KO mice, we reasoned that Arc expression each day at the time of testing may be important, thus we examined protein levels 24 hours after the last IVSA session. We observed after cocaine IVSA that *Fmr1* KO mice have normal total homogenate (S1) levels of Arc in both NAc and DS but decreased synaptic expression compared to WT mice, which reached statistical significance for the NAc. This finding suggests that striatal Arc expression and/or synaptic localization may be destabilized by loss of FMRP and is particularly intriguing considering contrasting observations elsewhere in the brain. For example, when using whole-brain preparations, the percent of messenger RNA on polyribosomes and percent protein for Arc is increased in *Fmr1* KO mice, an effect further exaggerated in synaptoneurosome fractions (Zalfa *et al.*, 2003). Our results following drug exposure suggest the possibility that FMRP differentially regulates Arc in striatal cells, and also highlights the possibility that FMRP and Arc work in the same pathway to facilitate drug-related behavioral plasticity, though additional studies are needed to test these questions.

Lastly, impairments have been observed for *Fmr1* KO and conditional KO mice in flexibility, learning, and memory, including tasks motivated by either reward or aversion (D'Hooge *et al.*, 1997; Mineur *et al.*, 2002; Ding *et al.*, 2014; Nolan *et al.*, 2017), and those explicitly requiring hippocampal function (Zhao *et al.*, 2005; Guo *et al.*, 2011; Arbab *et al.*, 2018). In general, deficits appear to be significantly milder on the C57BL/6 background strain (Spencer *et al.*, 2011). Here, we find that *Fmr1* KO (C57BL/6N) mice normally acquire and extinguish operant self-administration with either cocaine or a palatable liquid food as the reinforcer, suggesting that FMRP is not required for instrumental learning. Previous studies using natural reinforcers have reported normal acquisition of operant-based tasks in *Fmr1* KO (C57BL/6) mice, including olfactory discrimination (Larson *et al.*, 2008), but at least one found exaggerated extinction in *Fmr1* KO mice using an appetitive, two-choice operant task (Sidorov *et al.*, 2014). One possible reason for our different observation in food operant extinction is our use of vehicle (water) during this phase. In any case, our results add to the literature distinguishing roles for FMRP in different types of learning and suggest that FMRP’s role in facilitating adjustments in early reward-related behavior may be specific to drug reward.

## Conclusions

Here, we report that *Fmr1* KO mice have a significantly downward-shifted cocaine IVSA dose-response curve compared to WT counterparts, and they later decrease responding for cocaine at the acquisition dose over increasing schedules of reinforcement. Overall, our findings reveal a critical role for FMRP in mediating behavioral plasticity during cocaine self-administration, in a manner independent of learning, a role which may facilitate shifts in hedonic set point following drug exposure and/or vulnerability to drug-taking at lower doses. We also identify decreased NAc synaptosome expression of the FMRP target, Arc, in *Fmr1* KO mice following cocaine IVSA. Alongside the similar behavioral results in both KO mouse lines, our findings suggest that FMRP and Arc may work together to facilitate cocaine IVSA behavioral changes, though additional studies are required. Moving forward, it will also be important to investigate in what cell types, and the extent to which, FMRP facilitates cocaine-related behaviors, including in the development of compulsive drug-seeking and reinstatement, key features of substance use disorders.

## Acknowledgments

We thank Dr. Morgane Thomsen for consultation on mouse intravenous self-administration, and Dorothy Foster for behavioral testing assistance. This work was supported by the NIDA Drug Supply Program (gifted drug) and Texas A&M University (L.N.S.).

## Author Contributions

J.H. contributed to the design, acquisition, analysis, interpretation, and writing of the manuscript. M.C.D. contributed to the design, acquisition, analysis, and revision of this manuscript. C.T.P. contributed to the acquisition and revision of the manuscript. Y.G. contributed to the design, acquisition, and revision of the manuscript. L.N.S. contributed to the design, acquisition, analysis, interpretation, and writing of the manuscript.

## Abbreviations

DRD1: dopamine receptor D1
FMRP: fragile X mental retardation protein
FR: fixed ratio
FXS: fragile X syndrome
IVSA: intravenous self-administration
KO: knockout
MV: multivariate
SME: simple main effect
WT: wild type

