## Supplementary material for "The fragile X mental retardation protein promotes adjustments in cocaine self-administration that preserve reinforcement level": Figs. S1, S2

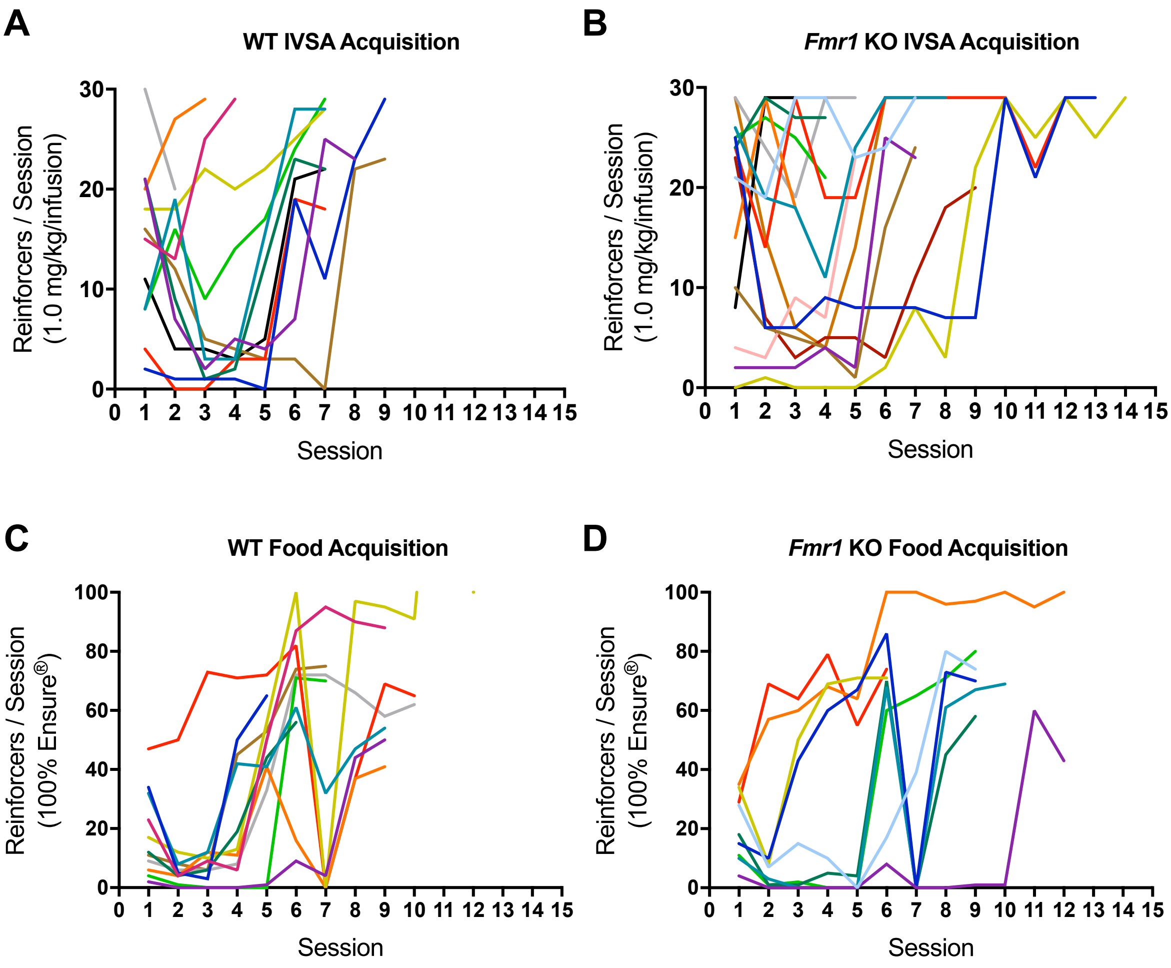


**Figure S1.** Individual WT (A, C) and *Fmr1* KO (B, D) active port, reinforced nose-pokes by session during cocaine IVSA (1.0 mg/kg/infusion) and food operant conditioning (20 ul 100% vanilla-flavored Ensure® each) acquisition, respectively.

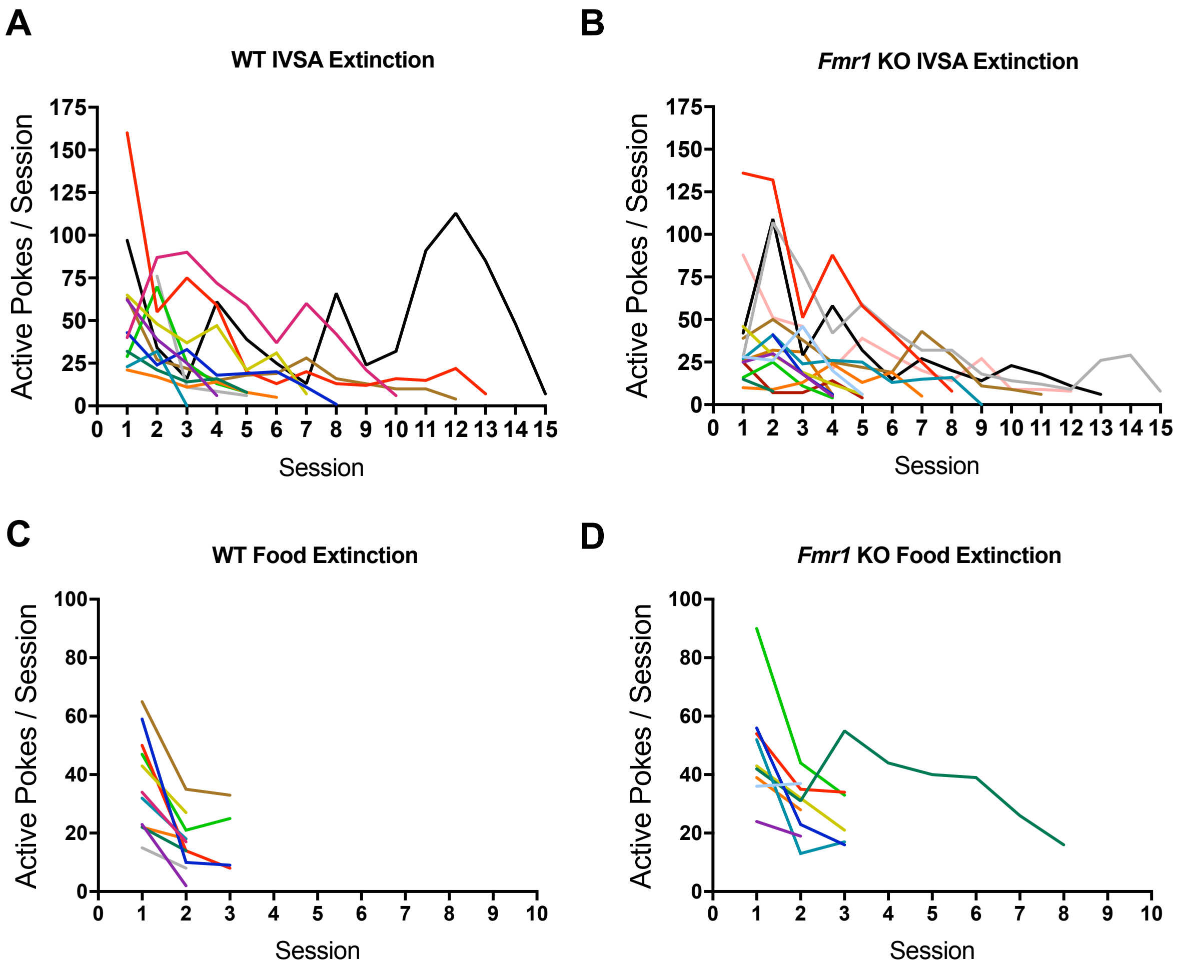


**Figure S2.** Individual WT (A, C) and *Fmr1* KO (B, D) active port (non-reinforced) nose-pokes by session during cocaine IVSA and food operant conditioning extinction, respectively.
