## Supplementary material for "The fragile X mental retardation protein promotes adjustments in cocaine self-administration that preserve reinforcement level": Table S1

| Figure | Statistical Analysis | Dependent Variable | Factor(s) | DF | *F* / *t* / $X^2$ value | *P* value |
|---|---|---|---|---|---|---|
| 2B | Kaplan-Meier log rank (Mantel-Cox) | group survival distribution to reaching acquisition criteria (drug) | genotype | 1 | 1.718 | 0.19 |
| 2C | Three-Way RM ANOVA | active vs inactive port, first and last day of acquisition (drug) | genotype | 1, 24 | 0.035 | 0.853 |
| | | | session | 1, 24 | 2.044 | 0.166 |
| | | | port | 1, 24 | 46.521 | < 0.0001 |
| | | | genotype x session | 1, 24 | 0.371 | 0.548 |
| | | | genotype x port | 1, 24 | 0.806 | 0.378 |
| | | | **session x port** | **1, 24** | **72.277** | **< 0.0001** |
| | | | genotype x session x port | 1, 24 | 3.112 | 0.09 |
| | One-Way RM ANOVA | active vs inactive port nosepokes, first day of acquisition (drug) | port | 1, 25 | 0.796 | 0.381 |
| | One-Way RM ANOVA | active vs inactive port nosepokes, last day of acquisition (drug) | **port** | **1, 26** | **499.497** | **< 0.0001** |
| | One-Way RM ANOVA | first vs last acquisition session, active port (drug) | **session** | **1, 26** | **23.159** | **< 0.0001** |
| | One-Way RM ANOVA | first vs last acquisition session, inactive port (drug) | **session** | **1, 25** | **25.915** | **< 0.0001** |
| 2E | Kaplan-Meier log rank (Mantel-Cox) | group survival distribution to reaching acquisition criteria (food) | genotype | 1 | 0.0622 | 0.8031 |
| 2F | Three-Way RM ANOVA | active vs inactive port, first and last day of acquisition (food) | genotype | 1, 18 | 0.319 | 0.579 |
| | | | session | 1, 18 | 21.927 | 0.000185 |
| | | | port | 1, 18 | 96.502 | < 0.0001 |
| | | | genotype x session | 1, 18 | 0.077 | 0.785 |
| | | | genotype x port | 1, 18 | 0.199 | 0.661 |
| | | | **session x port** | **1, 18** | **104.585** | **< 0.0001** |
| | | | genotype x session x port | 1, 18 | 0.598 | 0.449 |
| | One-Way RM ANOVA | active vs inactive port nosepokes, first day of acquisition (food) | **port** | **1, 19** | **5.438** | **0.031** |
| | One-Way RM ANOVA | active vs inactive port nosepokes, last day of acquisition (food) | **port** | **1, 19** | **206.64** | **< 0.0001** |
| | One-Way RM ANOVA | first vs last acquisition session, active port (food) | **session** | **1, 19** | **172.929** | **< 0.0001** |
| | One-Way RM ANOVA | first vs last acquisition session, inactive port (food) | session | 1, 19 | 4.279 | 0.052 |
| **Figure** | **Statistical Analysis** | **Dependent Variable** | **Factor(s)** | **DF** | ***F* / *t* / $X^2$ value** | ***P* value** |
| 3B | Kaplan-Meier log rank (Mantel-Cox) | group survival distribution to reaching extinction criteria (drug) | genotype | 1 | 0.4344 | 0.5099 |
| 3C | Three-Way RM ANOVA | active vs inactive port, first and last day of extinction (drug) | genotype | 1, 21 | 0.718 | 0.406 |
| | | | session | 1, 21 | 54.459 | < 0.0001 |
| | | | port | 1, 21 | 15.107 | 0.001 |
| | | | genotype x session | 1, 21 | 1.345 | 0.259 |
| | | | genotype x port | 1, 21 | 2.497 | 0.129 |
| | | | **session x port** | **1, 21** | **22.462** | **< 0.0001** |
| | | | genotype x session x port | 1, 21 | 2.972 | 0.099 |
| | One-Way RM ANOVA | active vs inactive port nosepokes, first day of extinction (drug) | **port** | **1, 22** | **16.352** | **0.001** |
| | One-Way RM ANOVA | active vs inactive port nosepokes, last day of extinction (drug) | port | 1, 26 | 0.126 | 0.726 |
| | One-Way RM ANOVA | first vs last extinction session, active port (drug) | **session** | **1, 22** | **52.214** | **< 0.0001** |
| | One-Way RM ANOVA | first vs last extinction session, inactive port (drug) | **session** | **1, 25** | **10.466** | **0.003** |
| 3E | Kaplan-Meier log rank (Mantel-Cox) | group survival distribution to reaching extinction criteria (food) | genotype | 1 | 2.34 | 0.1261 |
| 3F | Three-Way RM ANOVA | active vs inactive port, first and last day of extinction (food) | genotype | 1, 17 | 1.39 | 0.255 |
| | | | session | 1, 17 | 20.68 | 0.000285 |
| | | | port | 1, 17 | 43.489 | 0.000005 |
| | | | genotype x session | 1, 17 | 0.225 | 0.641 |
| | | | genotype x port | 1, 17 | 2.065 | 0.169 |
| | | | **session x port** | **1, 17** | **37.79** | **< 0.0001** |
| | | | genotype x session x port | 1, 17 | 2.011 | 0.174 |
| | One-Way RM ANOVA | active vs inactive port nosepokes, first day of extinction (food) | **port** | **1, 18** | **49.484** | **< 0.0001** |
| | One-Way RM ANOVA | active vs inactive port nosepokes, last day of extinction (food) | **port** | **1, 19** | **11.978** | **0.003** |
| | One-Way RM ANOVA | first vs last extinction session, active port (food) | **session** | **1, 18** | **40.236** | **< 0.0001** |
| | One-Way RM ANOVA | first vs last extinction session, inactive port (food) | session | 1, 18 | 2.718 | 0.117 |
| **Figure** | **Statistical Analysis** | **Dependent Variable** | **Factor(s)** | **DF** | ***F* / *t* / $X^2$ value** | ***P* value** |
| 4A | Three-Way RM ANOVA | active vs inactive port nosepokes per session for DR (drug) | genotype | 1, 17 | 4.711 | 0.044 |
| | | | dose | 2.1, 17 | 11.349 | < 0.0001 |
| | | | port | 1, 17 | 47.856 | < 0.0001 |
| | | | genotype x dose | 2.1, 17 | 3.006 | 0.059 |
| | | | genotype x port | 1, 17 | 3.707 | 0.071 |
| | | | dose x port | 2.2, 36.8 | 14.615 | < 0.0001 |
| | | | **genotype x dose x port** | **2.2, 36.8** | **3.235** | **0.047** |
| | Two-Way RM ANOVA | WT vs KO active port nosepokes, DR (drug) | genotype | 1, 19 | 6.529 | 0.019 |
| | | | dose | 2.1, 39.6 | 17.949 | < 0.0001 |
| | | | **genotype x dose** | **2.1, 39.6** | **4.873** | **0.012** |
| | One-Way RM ANOVA | active port nosepokes over dose, WT only (drug) | **dose** | **2.1, 16.6** | **14.902** | **< 0.0001** |
| | Bonferroni Pairwise Comparison | WT active port nosepokes, 0.01 vs. 0.032 mg/kg/infusion (drug) | dose | | | 1 |
| | Bonferroni Pairwise Comparison | WT active port nosepokes, 0.01 vs. 0.1 mg/kg/infusion (drug) | dose | | | 0.544 |
| | Bonferroni Pairwise Comparison | WT active port nosepokes, 0.01 vs. 0.32 mg/kg/infusion (drug) | **dose** | | | **0.01** |
| | Bonferroni Pairwise Comparison | WT active port nosepokes, 0.01 vs. 1.0 mg/kg/infusion (drug) | dose | | | 1 |
| | Bonferroni Pairwise Comparison | WT active port nosepokes, 0.01 vs. 3.2 mg/kg/infusion (drug) | dose | | | 0.173 |
| | Bonferroni Pairwise Comparison | WT active port nosepokes, 0.032 vs. 0.1 mg/kg/infusion (drug) | dose | | | 0.336 |
| | Bonferroni Pairwise Comparison | WT active port nosepokes, 0.032 vs. 0.32 mg/kg/infusion (drug) | **dose** | | | **0.014** |
| | Bonferroni Pairwise Comparison | WT active port nosepokes, 0.032 vs. 1.0 mg/kg/infusion (drug) | dose | | | 1 |
| | Bonferroni Pairwise Comparison | WT active port nosepokes, 0.032 vs. 3.2 mg/kg/infusion (drug) | dose | | | 0.204 |
| | Bonferroni Pairwise Comparison | WT active port nosepokes, 0.1 vs. 0.32 mg/kg/infusion (drug) | dose | | | 1 |
| | Bonferroni Pairwise Comparison | WT active port nosepokes, 0.1 vs. 1.0 mg/kg/infusion (drug) | dose | | | 1 |
| | Bonferroni Pairwise Comparison | WT active port nosepokes, 0.1 vs. 3.2 mg/kg/infusion (drug) | dose | | | 0.172 |
| | Bonferroni Pairwise Comparison | WT active port nosepokes, 0.32 vs. 1.0 mg/kg/infusion (drug) | **dose** | | | **0.009** |
| | Bonferroni Pairwise Comparison | WT active port nosepokes, 0.32 vs. 3.2 mg/kg/infusion (drug) | **dose** | | | **0.002** |
| | Bonferroni Pairwise Comparison | WT active port nosepokes, 1.0 vs 3.2 mg/kg/infusion (drug) | **dose** | | | **< 0.0001** |
| | One-Way RM ANOVA | active port nosepokes over dose, KO only (drug) | **dose** | **2.0, 21.9** | **4.088** | **0.031** |
| | Bonferroni Pairwise Comparison | KO active port nosepokes, 0.01 vs. 0.032 mg/kg/infusion (drug) | dose | | | 1 |
| | Bonferroni Pairwise Comparison | KO active port nosepokes, 0.01 vs. 0.1 mg/kg/infusion (drug) | dose | | | 0.898 |
| | Bonferroni Pairwise Comparison | KO active port nosepokes, 0.01 vs. 0.32 mg/kg/infusion (drug) | dose | | | 0.871 |
| | Bonferroni Pairwise Comparison | KO active port nosepokes, 0.01 vs. 1.0 mg/kg/infusion (drug) | dose | | | 0.119 |
| | Bonferroni Pairwise Comparison | KO active port nosepokes, 0.01 vs. 3.2 mg/kg/infusion (drug) | dose | | | 0.485 |
| | Bonferroni Pairwise Comparison | KO active port nosepokes, 0.032 vs. 0.1 mg/kg/infusion (drug) | dose | | | 1 |
| | Bonferroni Pairwise Comparison | KO active port nosepokes, 0.032 vs. 0.32 mg/kg/infusion (drug) | dose | | | 1 |
| | Bonferroni Pairwise Comparison | KO active port nosepokes, 0.032 vs. 1.0 mg/kg/infusion (drug) | dose | | | 0.565 |
| | Bonferroni Pairwise Comparison | KO active port nosepokes, 0.032 vs. 3.2 mg/kg/infusion (drug) | dose | | | 0.153 |
| | Bonferroni Pairwise Comparison | KO active port nosepokes, 0.1 vs. 0.32 mg/kg/infusion (drug) | dose | | | 1 |
| | Bonferroni Pairwise Comparison | KO active port nosepokes, 0.1 vs. 1.0 mg/kg/infusion (drug) | dose | | | 1 |
| | Bonferroni Pairwise Comparison | KO active port nosepokes, 0.1 vs. 3.2 mg/kg/infusion (drug) | dose | | | 0.262 |
| | Bonferroni Pairwise Comparison | KO active port nosepokes, 0.32 vs. 1.0 mg/kg/infusion (drug) | dose | | | 1 |
| | Bonferroni Pairwise Comparison | KO active port nosepokes, 0.32 vs. 3.2 mg/kg/infusion (drug) | dose | | | 0.352 |
| | Bonferroni Pairwise Comparison | KO active port nosepokes, 1.0 vs 3.2 mg/kg/infusion (drug) | **dose** | | | **< 0.0001** |
| | One-way MANOVA | WT vs KO active port nosepokes, 0.01 mg/kg/infusion (drug) | genotype | 1, 17 | 3.614 | 0.074 |
| | One-way MANOVA | WT vs KO active port nosepokes, 0.032 mg/kg/infusion (drug) | genotype | 1, 17 | 0.907 | 0.354 |
| | One-way MANOVA | WT vs KO active port nosepokes, 0.1 mg/kg/infusion (drug) | genotype | 1, 17 | 1.175 | 0.294 |
| | One-way MANOVA | WT vs KO active port nosepokes, 0.32 mg/kg/infusion (drug) | **genotype** | **1, 17** | **8.023** | **0.011** |
| | One-way MANOVA | WT vs KO active port nosepokes, 1.0 mg/kg/infusion (drug) | genotype | 1, 17 | 0.719 | 0.408 |
| | One-way MANOVA | WT vs KO active port nosepokes, 3.2 mg/kg/infusion (drug) | genotype | 1, 17 | 0.897 | 0.357 |

| Figure | Statistical Analysis | Dependent Variable | Factor(s) | DF | $F$ / $t$ / $X^2$ value | $P$ value |
|---|---|---|---|---|---|---|
| | Two-Way RM ANOVA | WT vs KO inactive port nosepokes, DR (drug) | genotype | 1, 17 | 0.803 | 0.383 |
| | | | dose | 4.3, 73.7 | 1.118 | 0.356 |
| | | | genotype x dose | 4.3, 73.7 | 1.432 | 0.229 |
| | Two-Way RM ANOVA | active vs inactive port nosepokes, DR, WT only (drug) | port | 1, 6 | 25.738 | 0.002 |
| | | | dose | 1.8, 11 | 7.469 | 0.01 |
| | | | **port x dose** | **2.1, 12.7** | **9.636** | **0.003** |
| | One-Way RM ANOVA | active vs inactive port nosepokes, 0.01 mg/kg/infusion, WT only (drug) | port | 1, 8 | 1.121 | 0.321 |
| | One-Way RM ANOVA | active vs inactive port nosepokes, 0.032 mg/kg/infusion, WT only (drug) | port | 1, 7 | 4.933 | 0.062 |
| | One-Way RM ANOVA | active vs inactive port nosepokes, 0.1 mg/kg/infusion, WT only (drug) | **port** | **1, 6** | **8.677** | **0.026** |
| | One-Way RM ANOVA | active vs inactive port nosepokes, 0.32 mg/kg/infusion, WT only (drug) | **port** | **1, 8** | **46.076** | **0.00014** |
| | One-Way RM ANOVA | active vs inactive port nosepokes, 1.0 mg/kg/infusion, WT only (drug) | **port** | **1, 8** | **93.392** | **< 0.0001** |
| | One-Way RM ANOVA | active vs inactive port nosepokes, 3.2 mg/kg/infusion, WT only (drug) | port | 1, 8 | 0.339 | 0.576 |
| | Two-Way RM ANOVA | active vs inactive port nosepokes, DR, KO only (drug) | port | 1, 11 | 19.016 | 0.001 |
| | | | dose | 2.2, 24.5 | 3.393 | 0.046 |
| | | | **port x dose** | **2.0, 22.4** | **4.703** | **0.019** |
| | One-Way RM ANOVA | active vs inactive port nosepokes, 0.01 mg/kg/infusion, KO only (drug) | **port** | **1, 11** | **11.462** | **0.006** |
| | One-Way RM ANOVA | active vs inactive port nosepokes, 0.032 mg/kg/infusion, KO only (drug) | port | 1, 11 | 4.81 | 0.051 |
| | One-Way RM ANOVA | active vs inactive port nosepokes, 0.1 mg/kg/infusion, KO only (drug) | **port** | **1, 11** | **8.939** | **0.012** |
| | One-Way RM ANOVA | active vs inactive port nosepokes, 0.32 mg/kg/infusion, KO only (drug) | **port** | **1, 11** | **8.84** | **0.013** |
| | One-Way RM ANOVA | active vs inactive port nosepokes, 1.0 mg/kg/infusion, KO only (drug) | **port** | **1, 11** | **50.557** | **< 0.0001** |
| | One-Way RM ANOVA | active vs inactive port nosepokes, 3.2 mg/kg/infusion, KO only (drug) | port | 1, 11 | 2.908 | 0.116 |
| 4B | Two-Way RM ANOVA | total drug intake, dose response (drug) | genotype | 1, 19 | 7.021 | 0.016 |
| | | | dose | 2.7, 52.2 | 79.506 | < 0.0001 |
| | | | **genotype x dose** | **2.7, 52.2** | **5.374** | **0.003** |
| | One-way MANOVA | WT vs KO, total drug intake, 0.01 mg/kg/infusion (drug) | genotype | 1, 19 | 2.782 | 0.112 |
| | One-way MANOVA | WT vs KO, total drug intake, 0.032 mg/kg/infusion (drug) | genotype | 1, 19 | 0.894 | 0.356 |
| | One-way MANOVA | WT vs KO, total drug intake, 0.1 mg/kg/infusion (drug) | genotype | 1, 19 | 1.428 | 0.247 |
| | One-way MANOVA | WT vs KO, total drug intake, 0.32 mg/kg/infusion (drug) | **genotype** | **1, 19** | **12.526** | **0.002** |
| | One-way MANOVA | WT vs KO, total drug intake, 1.0 mg/kg/infusion (drug) | genotype | 1, 19 | 1.282 | 0.272 |
| | One-way MANOVA | WT vs KO, total drug intake, 3.2 mg/kg/infusion (drug) | genotype | 1, 19 | 2.024 | 0.171 |
| | One-way RM ANOVA | total drug intake across doses, WT only (drug) | **dose** | **2.2, 17.5** | **60.58** | **< 0.0001** |
| | Bonferroni Pairwise Comparison | WT drug intake, 0.01 vs 0.032 mg/kg/infusion (drug) | dose | | | 0.051 |
| | Bonferroni Pairwise Comparison | WT drug intake, 0.01 vs 0.1 mg/kg/infusion (drug) | dose | | | 0.09 |
| | Bonferroni Pairwise Comparison | WT drug intake, 0.01 vs 0.32 mg/kg/infusion (drug) | **dose** | | | **0.001** |
| | Bonferroni Pairwise Comparison | WT drug intake, 0.01 vs 1.0 mg/kg/infusion (drug) | **dose** | | | **< 0.0001** |
| | Bonferroni Pairwise Comparison | WT drug intake, 0.01 vs 3.2 mg/kg/infusion (drug) | **dose** | | | **< 0.0001** |
| | Bonferroni Pairwise Comparison | WT drug intake, 0.032 vs 0.1 mg/kg/infusion (drug) | dose | | | 0.112 |
| | Bonferroni Pairwise Comparison | WT drug intake, 0.032 vs 0.32 mg/kg/infusion (drug) | **dose** | | | **0.002** |
| | Bonferroni Pairwise Comparison | WT drug intake, 0.032 vs 1.0 mg/kg/infusion (drug) | **dose** | | | **< 0.0001** |
| | Bonferroni Pairwise Comparison | WT drug intake, 0.032 vs 3.2 mg/kg/infusion (drug) | **dose** | | | **< 0.0001** |
| | Bonferroni Pairwise Comparison | WT drug intake, 0.1 vs 0.32 mg/kg/infusion (drug) | **dose** | | | **0.003** |
| | Bonferroni Pairwise Comparison | WT drug intake, 0.1 vs 1.0 mg/kg/infusion (drug) | **dose** | | | **< 0.0001** |
| | Bonferroni Pairwise Comparison | WT drug intake, 0.1 vs 3.2 mg/kg/infusion (drug) | **dose** | | | **0.001** |
| | Bonferroni Pairwise Comparison | WT drug intake, 0.32 vs 1.0 mg/kg/infusion (drug) | dose | | | 1 |
| | Bonferroni Pairwise Comparison | WT drug intake, 0.32 vs 3.2 mg/kg/infusion (drug) | dose | | | 1 |
| | Bonferroni Pairwise Comparison | WT drug intake, 1.0 vs 3.2 mg/kg/infusion (drug) | dose | | | 1 |
| | One-way RM ANOVA | total drug intake across doses, KO only (drug) | **dose** | **2.5, 27.4** | **31.402** | **< 0.0001** |
| | Bonferroni Pairwise Comparison | KO drug intake, 0.01 vs 0.032 mg/kg/infusion (drug) | **dose** | | | **0.016** |
| | Bonferroni Pairwise Comparison | KO drug intake, 0.01 vs 0.1 mg/kg/infusion (drug) | dose | | | 0.108 |
| | Bonferroni Pairwise Comparison | KO drug intake, 0.01 vs 0.32 mg/kg/infusion (drug) | dose | | | 0.142 |
| | Bonferroni Pairwise Comparison | KO drug intake, 0.01 vs 1.0 mg/kg/infusion (drug) | **dose** | | | **< 0.0001** |
| | Bonferroni Pairwise Comparison | KO drug intake, 0.01 vs 3.2 mg/kg/infusion (drug) | **dose** | | | **0.002** |
| | Bonferroni Pairwise Comparison | KO drug intake, 0.032 vs 0.1 mg/kg/infusion (drug) | dose | | | 0.203 |
| | Bonferroni Pairwise Comparison | KO drug intake, 0.032 vs 0.32 mg/kg/infusion (drug) | dose | | | 0.161 |
| | Bonferroni Pairwise Comparison | KO drug intake, 0.032 vs 1.0 mg/kg/infusion (drug) | **dose** | | | **< 0.0001** |
| | Bonferroni Pairwise Comparison | KO drug intake, 0.032 vs 3.2 mg/kg/infusion (drug) | **dose** | | | **0.002** |
| | Bonferroni Pairwise Comparison | KO drug intake, 0.1 vs 0.32 mg/kg/infusion (drug) | dose | | | 0.632 |
| | Bonferroni Pairwise Comparison | KO drug intake, 0.1 vs 1.0 mg/kg/infusion (drug) | **dose** | | | **< 0.0001** |
| | Bonferroni Pairwise Comparison | KO drug intake, 0.1 vs 3.2 mg/kg/infusion (drug) | **dose** | | | **0.008** |
| | Bonferroni Pairwise Comparison | KO drug intake, 0.32 vs 1.0 mg/kg/infusion (drug) | **dose** | | | **0.004** |
| | Bonferroni Pairwise Comparison | KO drug intake, 0.32 vs 3.2 mg/kg/infusion (drug) | dose | | | 0.058 |
| | Bonferroni Pairwise Comparison | KO drug intake, 1.0 vs 3.2 mg/kg/infusion (drug) | dose | | | 1 |
| 4C | Three-Way RM ANOVA | active vs inactive port nosepokes per session for DR (food) | genotype | 1, 18 | 1.141 | 0.299 |
| | | | concentration | 3, 54 | 54.106 | < 0.0001 |
| | | | port | 1, 18 | 358.199 | < 0.0001 |
| | | | genotype x concentration | 3, 54 | 0.698 | 0.557 |
| | | | genotype x port | 1, 18 | 1.669 | 0.213 |
| | | | **concentration x port** | **3, 54** | **79.627** | **< 0.0001** |
| | | | genotype x concentration x port | 3, 54 | 0.731 | 0.538 |
| | Two-Way R:M ANOVA | WT vs KO, active nosepokes, DR (food) | genotype | 1, 18 | 1.398 | 0.252 |
| | | | **concentration** | **3, 54** | **73.575** | **< 0.0001** |
| | | | genotype x concentration | 3, 54 | 0.616 | 0.607 |
| | Two-Way R:M ANOVA | WT vs KO, inactive nosepokes, DR (food) | genotype | 1, 18 | 0.38 | 0.545 |
| | | | **concentration** | **3, 54** | **3.159** | **0.032** |
| | | | genotype x concentration | 3, 54 | 1.337 | 0.272 |
| | Two-Way RM ANOVA | active vs inactive port nosepokes, DR, WT only (food) | port | 1, 10 | 176.82 | < 0.0001 |
| | | | concentration | 3, 30 | 33.739 | < 0.0001 |
| | | | **port x concentration** | **3, 30** | **40.546** | **< 0.0001** |
| | One-Way RM ANOVA | active port nosepokes over concentrations, WT only (food) | **concentration** | **3, 30** | **42.305** | **< 0.0001** |
| | Bonferroni Pairwise Comparison | WT active port nosepokes, 3 vs 10% (food) | **concentration** | | | **0.005** |
| | Bonferroni Pairwise Comparison | WT active port nosepokes, 3 vs 32% (food) | **concentration** | | | **< 0.0001** |
| | Bonferroni Pairwise Comparison | WT active port nosepokes, 3 vs 100% (food) | **concentration** | | | **0.000002** |
| | Bonferroni Pairwise Comparison | WT active port nosepokes, 10% vs 32% (food) | **concentration** | | | **0.007** |
| | Bonferroni Pairwise Comparison | WT active port nosepokes, 10% vs 100% (food) | concentration | | | 0.966 |
| | Bonferroni Pairwise Comparison | WT active port nosepokes, 32% vs 100% (food) | **concentration** | | | **0.006** |
| | One-Way RM ANOVA | inactive port nosepokes over concentrations, WT only (food) | concentration | 3, 30 | 0.537 | 0.66 |
| | Two-Way RM ANOVA | active vs inactive port nosepokes, DR, KO only (food) | port | 1, 8 | 180.618 | < 0.0001 |
| | | | concentration | 3, 24 | 22.23 | < 0.0001 |
| | | | **port x concentration** | **3, 24** | **45.959** | **< 0.0001** |
| | One-Way RM ANOVA | active port nosepokes over concentrations, KO only (food) | **concentration** | **3, 24** | **33.915** | **< 0.0001** |
| | Bonferroni Pairwise Comparison | KO active port nosepokes, 3 vs 10% (food) | **concentration** | | | **0.026** |
| | Bonferroni Pairwise Comparison | KO active port nosepokes, 3 vs 32% (food) | **concentration** | | | **< 0.0001** |
| | Bonferroni Pairwise Comparison | KO active port nosepokes, 3 vs 100% (food) | **concentration** | | | **0.00025073** |
| | Bonferroni Pairwise Comparison | KO active port nosepokes, 10% vs 32% (food) | **concentration** | | | **0.014** |
| | Bonferroni Pairwise Comparison | KO active port nosepokes, 10% vs 100% (food) | concentration | | | 1 |
| | Bonferroni Pairwise Comparison | KO active port nosepokes, 32% vs 100% (food) | **concentration** | | | **0.006** |
| | One-Way RM ANOVA | inactive port nosepokes over concentrations, KO only (food) | **concentration** | **3, 24** | **4.479** | **0.012** |
| | Bonferroni Pairwise Comparison | KO inactive port nosepokes, 3 vs 10% (food) | concentration | | | 1 |
| | Bonferroni Pairwise Comparison | KO inactive port nosepokes, 3 vs 32% (food) | concentration | | | 1 |

| Figure | Statistical Analysis | Dependent Variable | Factor(s) | DF | *F* / *t* / $X^2$ value | *P* value |
|---|---|---|---|---|---|---|
| | Bonferroni Pairwise Comparison | KO inactive port nosepokes, 3 vs 100% (food) | concentration | | | 0.528 |
| | Bonferroni Pairwise Comparison | KO inactive port nosepokes, 10% vs 32% (food) | concentration | | | 1 |
| | Bonferroni Pairwise Comparison | KO inactive port nosepokes, 10% vs 100% (food) | concentration | | | 0.051 |
| | Bonferroni Pairwise Comparison | KO inactive port nosepokes, 32% vs 100% (food) | concentration | | | 0.104 |
| 4D | Four-Way RM ANOVA | active vs inactive port nosepokes across increasing cost (drug) | **genotype** | **1, 9** | **5.795** | **0.039** |
| | | | cost | 1.7, 15.4 | 7.302 | 0.008 |
| | | | day | 1, 9 | 0.454 | 0.517 |
| | | | port | 1, 9 | 35.488 | 0.0002 |
| | | | genotype x cost | 1.7, 15.4 | 0.843 | 0.433 |
| | | | genotype x port | 1, 9 | 1.259 | 0.291 |
| | | | genotype x day | 1, 9 | 1.503 | 0.251 |
| | | | cost x day | 2, 18 | 0.139 | 0.871 |
| | | | **cost x port** | **2, 18** | **28.997** | **< 0.0001** |
| | | | day x port | 1, 9 | 0.117 | 0.74 |
| | | | genotype x cost x day | 2, 18 | 2.168 | 0.143 |
| | | | genotype x cost x port | 2, 18 | 1.561 | 0.237 |
| | | | genotype x day x port | 1, 9 | 0.223 | 0.648 |
| | | | port x cost x day | 2, 18 | 0.738 | 0.492 |
| | | | port x cost x day x genotype | 2, 18 | 2.218 | 0.138 |
| | One-Way RM ANOVA | active vs inactive pokes, FR1 | **port** | **1, 10** | **117.356** | **< 0.0001** |
| | One-Way RM ANOVA | active vs inactive pokes, FR3 | **port** | **1, 10** | **15.121** | **0.003** |
| | One-Way RM ANOVA | active vs inactive pokes, FR5 | port | 1, 10 | 4.291 | 0.065 |
| | One-Way RM ANOVA | active nosepokes over increasing cost | **cost** | **1.6, 15.9** | **25.741** | **< 0.0001** |
| | Bonferroni Pairwise Comparison | active nosepokes, FR1 vs FR3 | **cost** | | | **0.003** |
| | Bonferroni Pairwise Comparison | active nosepokes, FR1 vs FR5 | **cost** | | | **0.001** |
| | Bonferroni Pairwise Comparison | active nosepokes, FR3 vs FR5 | **cost** | | | **0.007** |
| | One-Way RM ANOVA | inactive nosepokes over increasing cost | cost | 1.2, 11.5 | 1.679 | 0.224 |
| | One-Way MANOVA | WT vs KO active port nosepokes at FR1 | genotype | 1, 10 | 3.946 | 0.078 |
| | One-Way MANOVA | WT vs KO active port nosepokes at FR3 | genotype | 1, 10 | 3.366 | 0.1 |
| | One-Way MANOVA | WT vs KO active port nosepokes at FR5 | **genotype** | **1, 10** | **5.739** | **0.04** |
| | One-Way MANOVA | WT vs KO inactive port nosepokes at FR1 | genotype | 1, 10 | 0.356 | 0.565 |
| | One-Way MANOVA | WT vs KO inactive port nosepokes at FR3 | genotype | 1, 10 | 2.022 | 0.189 |
| | One-Way MANOVA | WT vs KO inactive port nosepokes at FR5 | genotype | 1, 10 | 0.585 | 0.464 |

| Figure | Statistical Analysis | Dependent Variable | Factor(s) | DF | *F* / *t* / $X^2$ value | *P* value |
|---|---|---|---|---|---|---|
| 5B | Two-Way ANOVA | normalized PSD95/GAPDH, relative to WT S1 (NAc) | genotype | 1, 18 | 3.676 | 0.0712 |
| | | | **fraction** | **2, 18** | **40.17** | **< 0.0001** |
| | | | genotype x fraction | 2, 18 | 1.46 | 0.2585 |
| 5C | Two-Way ANOVA | normalized NR2B/GAPDH, relative to WT S1 (NAc) | genotype | 1, 18 | 1.048 | 0.3195 |
| | | | **fraction** | **2, 18** | **17.49** | **< 0.0001** |
| | | | genotype x fraction | 2, 18 | 0.01466 | 0.9855 |
| 5D | Two-Way ANOVA | normalized ARC/GAPDH, relative to WT S1 (NAc) | genotype | 1, 18 | 8.465 | 0.0094 |
| | | | fraction | 2, 18 | 31.4 | < 0.0001 |
| | | | **genotype x fraction** | **2, 18** | **6.228** | **0.0088** |
| | Bonferroni Post-hoc | S1 expression | genotype | 18 | 0.6065 | 1 |
| | Bonferroni Post-hoc | S2 expression | genotype | 18 | 0.9963 | 1 |
| | Bonferroni Post-hoc | P expression | **genotype** | **18** | **4.532** | **0.0039** |
| 5F | Two-Way ANOVA | normalized PSD95/GAPDH, relative to WT S1 (DS) | genotype | 1, 18 | 2.712 | 0.117 |
| | | | **fraction** | **2, 18** | **16.04** | **< 0.0001** |
| | | | genotype x fraction | 2, 18 | 1.532 | 0.2431 |
| 5G | Two-Way ANOVA | normalized NR2B/GAPDH, relative to WT S1 (DS) | genotype | 1, 18 | 1.057 | 0.3175 |
| | | | **fraction** | **2, 18** | **11.76** | **0.0005** |
| | | | genotype x fraction | 2, 18 | 1.761 | 0.2002 |
| 5H | Two-Way ANOVA | normalized ARC/GAPDH, relative to WT S1 (DS) | genotype | 1, 18 | 2.193 | 0.1559 |
| | | | **fraction** | **2, 18** | **10.31** | **0.001** |
| | | | genotype x fraction | 2, 18 | 2.09 | 0.1527 |
